# A novel 3-miRNA network regulates tumour progression in oral squamous cell carcinoma

**DOI:** 10.1101/2022.05.31.494114

**Authors:** Aditi Patel, Parina Patel, Dushyant Mandlik, Kaustubh Patel, Pooja Malaviya, Kaid Johar, Krishna B.S Swamy, Shanaya Patel, Vivek Tanavde

## Abstract

Oral squamous cell carcinoma (OSCC) is often diagnosed late, leading to poor patient outcomes. This study aims to identify potential miRNA-based biomarkers for predicting disease progression using salivary exosomes derived from OSCC patients. Further, we identify crucial miRNA-mRNA networks involved in tumorigenesis and uncover the underlying mechanism responsible for OSCC progression.

Small RNA (n=23) sequencing analysis along with data available from The Cancer Genome Atlas (TCGA) (n=114) identified 12 differentially expressed miRNAs in OSCC patients as compared to controls. Validating these findings, miR-140-5p, miR-143-5p, and miR-145-5p were significantly downregulated in a larger cohort of OSCC patients (n=70). This 3-miRNA signature demonstrated higher efficacy of salivary exosomes (p<0.0001) in early detection and clinically correlated with disease progression and overall survival of OSCC patients (p<0.05). Further, analysis of the transcriptome, TCGA datasets and miRNA-mRNA networks, identified top hub genes (HIF*1a*, *CDH1*, *CD44*, *EGFR*, and *CCND1*) which were regulated by a 3-miRNA signature. Based on pathway analysis, these miRNA-mRNA interactions were found to be involved in regulating epithelial-mesenchymal transition (EMT). Further, transfection-mediated upregulation of the 3-miRNA signature significantly decreased cell proliferation, induced apoptosis, resulted in G2/M phase cell cycle arrest and reduced the invasive and migratory potential by reversing the EMT process in OECM-1 cell line.

Thus, this study identifies a 3-miRNA signature that can be utilized as a potential biomarker for early detection of OSCC and uncovers the underlying mechanisms responsible for converting a normal epithelial cell into a malignant phenotype.

## Introduction

Oral squamous cell carcinoma (OSCC) is one of the major causes of morbidity and mortality worldwide (1). For over four decades, OSCC has been associated with an extremely low five-year survival rate, mainly due to late-stage detection. Despite recent advances in molecular diagnostics, no disease-specific biomarkers are clinically available for early risk prediction of OSCC. Conventional tissue biopsies fail to access deeper tumour sites and are unable to detect tumour heterogeneity (2, 3). Therefore, it is important to identify non-invasive biomarkers to facilitate the early diagnosis of OSCC.

Exosomal miRNAs have demonstrated immense clinical potential in early risk prediction, assessment of disease progression, and real-time therapeutic monitoring in various malignancies (4). Since miRNAs are highly tissue-specific, and negatively regulate 60% of protein-coding genes in humans, it is necessary to identify miRNA-based biomarkers and elucidate miRNA-mRNA networks that lead to the deregulation of various biological mechanisms and signalling pathways (5–7). So far, miRNA-mRNA network studies have failed to conduct a comprehensive analysis of small RNA and transcriptome profiles using matched samples. Moreover, these studies involved smaller patient cohorts and lacked functional validation using *in-vitro* or *in-vivo* models.

Thus, in this study, we conducted a comparative analysis of miRNA and transcriptome profiles of tumour tissues and salivary exosomes derived from OSCC patients. Further, we identified regulatory networks and underlying mechanisms responsible for tumour progression in OSCC. To the best of our knowledge, this is the first study to explore the potential clinical utility of saliva and tumour-derived miRNA-mRNA regulatory networks in OSCC.

## Methods

### Patient group

Resected tumour tissue and unstimulated saliva samples were collected from patients diagnosed with OSCC (excluding tongue, larynx, pharynx, and hypopharynx) at the HCG Cancer Centre, Ahmedabad. Patients were authorized for primary surgical resection by the institutional team after their malignancies were confirmed by histopathological examination. HPV, HIV, HCV and HbsAg-positive malignancies, pediatric cases and patients initially treated with other adjuvant therapeutic modalities were excluded from the study. All patients provided written informed consent for their participation in the study. This study was reviewed and approved by the HCG Cancer Centre’s Ethics Committee (ECR/92/Inst/GJ/2013/RR-16) for human subject research and the use of human tissues. This study also complied with the guidelines set forth by the Declaration of Helsinki (2008).

### Tissue and saliva collection

Samples were collected during R0 resections of early and late-stage OSCC. Resected specimens were evaluated histopathologically and residual tumour samples were snap-frozen and stored at −80 °C for further processing. For normal controls, brush biopsy samples were taken from healthy individuals (age and sex-matched with patients) with no clinically detectable oral lesions.

Whole saliva was collected in sterile tubes from healthy individuals and OSCC patients prior to surgical resection, using a protocol described in a previous study (4). The saliva samples were kept at −80 °C until further use. A total of 23 patient-derived samples and 21 samples from age- and sex-matched healthy volunteers were sequenced. Sequencing results were validated by qRT-PCR using a validation cohort of patients. The details of the samples are given in **Table 1.**

**Table 1:**
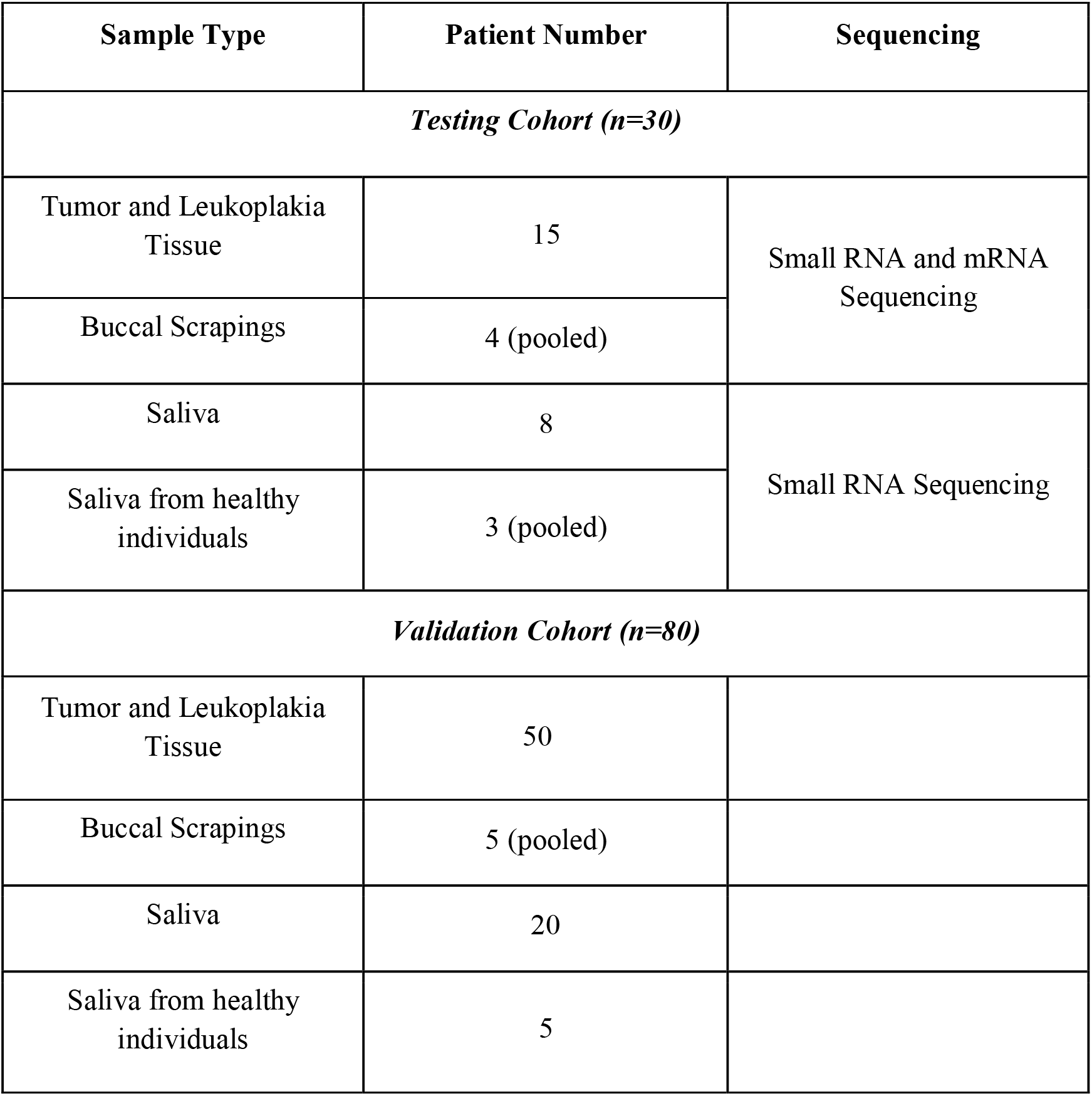
Sampling details.

| Sample Type | Patient Number | Sequencing |
| --- | --- | --- |
| Testing Cohort (n=30) |  |  |
| Tumor and Leukoplakia Tissue | 15 | Small RNA and mRNA Sequencing |
| Buccal Scrapings | 4 (pooled) |  |
| Saliva | 8 | Small RNA Sequencing |
| Saliva from healthy individuals | 3 (pooled) |  |
| Validation Cohort (n=80) |  |  |
| Tumor and Leukoplakia Tissue | 50 |  |
| Buccal Scrapings | 5 (pooled) |  |
| Saliva | 20 |  |
| Saliva from healthy individuals | 5 |  |

### Exosome isolation

Saliva from OSCC patients and healthy controls was centrifuged at 2000 × *g* for 10 min at 37°C to remove cells and debris. The supernatant was transferred to a sterile tube, and exosomes were precipitated using a commercially available kit as per the manufacturer’s protocol (Invitrogen^™^ Total Exosome Isolation Reagent (from other body fluids), Thermo Fisher Scientific). The exosome pellet was either re-suspended in 50μL of phosphate-buffered saline (PBS) for characterization or in 750μL of TRIzol LS reagent (Thermo Fisher Scientific) for storage at −80 °C for subsequent RNA extraction.

### Exosome characterization

The concentration (particles/mL) and size of exosomes were measured using NanoSight LM10 Nanoparticle Tracking Analysis (NTA) software (Malvern Instruments Ltd.). The exosome size was also determined by transmission electron microscopy (TEM) using 300 mesh carbon formvar grids. 5μL of 2% glutaraldehyde was added to the grids which were then wick-dried, rinsed, and stained with 2% uranyl acetate. The grids were examined using JEM1400 Plus transmission electron microscope (Jeol) at 100 kV. Images were obtained via a chamber scope film camera. The isolated exosomes were stained with anti-human CD9 allophycocyanin (CD9-APC), anti-human CD63 Alexa Fluor 488 (CD63-Alexa 488), and anti-human CD81 phycoerythrin (CD81-PE antibodies (Thermo Fisher Scientific). The samples were then analysed using the FACS Calibur^™^ Flow Cytometer (Becton Dickinson Biosciences).

### RNA extraction

Total RNA was extracted from tissue and salivary exosomes of OSCC patients using the described method (8). The RNA pellet was resuspended in 32μL of nuclease-free water. RNA yield and purity were quantified using Agilent 2100 Bioanalyzer (Agilent Technologies) and Qubit 4 Fluorometer (Thermo Fisher Scientific).

### miRNA and transcriptome sequencing

Small RNA and transcriptome sequencing of tumour tissues and salivary exosomes derived from OSCC patients was conducted at Genotypic Technology Pvt. Ltd., Bengaluru, India. The libraries were prepared using QIAseq^®^ miRNA library kit protocol (QIAGEN) as per the manufacturer’s instructions. Quality control and read mapping was performed as mentioned in our previous study (8). miRNAs were considered expressed when detected with a miRDeep2 score ≥ 5, reads >15 nucleotides in length, quality score >30 reads/sample, RPM normalized expression value□≥□the first quantile of expression distribution, false discovery rate (FDR) of ≤ 0.05, log_2_ fold-change (log_2_FC) of > 2, and statistically significant abundance in each sample (adj p-value < 0.05). This script produced a raw count file that was used as input for the DESeq2 R/Bioconductor package for further analysis (Supplementary Fig. S1).

For transcriptome profiling, library preparation was performed using NEBNext^®^ Ultra^™^ II directional RNA library prep kit and sequencing was performed on Illumina HiSeq 2500 platform using 150 × 2 chemistry. Raw sequencing reads were subjected to quality assessment using FastQC (9), followed by cleaning with Cutadapt (10). The refined reads were compared with a reference genome (GRCh, version 38) using Subread aligner (8, 11). Differential expression analysis was conducted using DESeq2 (RRID: SCR_000154) (12). Transcripts with read counts ≥ 30 were included. miRNAs and/or genes were considered differentially expressed based on the criteria: log_2_FC > 2 and the adjusted p-value < 0.01.

### Reverse transcription and quantitative Real-time Polymerase Chain Reaction (qRT-PCR)

For small RNA expression analysis of miR-140, miR-143, miR-145, miR-30a, miR-21, miR-423, and miR-let7a, total RNA was reverse transcribed using the miScript-II RT kit (QIAGEN) according to the manufacturer’s instructions. For quantifying the expression of genes (E-cadherin and N-cadherin), reverse transcription of RNA extracted from OECM-1 cells transfected with miRNA mimics and SC control was carried out using the Quantitect Reverse Transcription Kit (QIAGEN) according to the manufacturer’s protocol. The expression of identified miRNAs and genes were validated using the miScript SYBR^®^ Green PCR kit (QIAGEN) on the QuantStudio 5 Real-Time PCR system (Applied Biosystems). U6 snRNA and β-actin were used as endogenous controls for miRNA and mRNA PCRs respectively. Relative expression of miRNAs and genes was calculated using the 2^-ΔΔCt^ method. The sequences of the primers used are specified in Supplementary Table S1.

### miRNA target prediction

Candidate miRNA-mRNA target relationships were predicted by at least one or more of the following target prediction algorithms extracted from miRDB, TargetScan 5.1, and miRWalk (13–15). Genes were considered differentially expressed if log_2_FC>2, and adj p-value was ≤ 0.05.

### Interaction networks and pathway analysis

Protein-protein interaction and regulatory networks of the differentially expressed genes were generated using Cytoscape (RRID: SCR_003032) (v3.6.1) and Ingenuity Pathways Analysis software (IPA) (QIAGEN Inc., https://digitalinsights.qiagen.com/IPA) respectively (16, 17). The statistically significant canonical pathway analysis was conducted using IPA. Z-score >+2 was defined as the threshold of significant activation and/or inhibition.

### Cell Culture and transfection

The human oral squamous cell carcinoma cell line, OECM-1, was obtained from Sigma Aldrich. Cells were maintained in Roswell Park Memorial Institute (RPMI) 1640 Medium (Thermo Fisher Scientific) containing 10% Fetal Bovine Serum (FBS) (HiMedia) at 37°C in a humid atmosphere with 5% CO2. Transfection was performed using miR-143-5p, miR-145-5p, and miR-140-5p mimics and corresponding scrambled control (SC) (Ambion) using Lipofectamine 2000 reagent (Invitrogen) according to the manufacturer’s protocol. The total concentration of miR-143/miR-145/miR-140 mimics combination was 100nM and the transfection efficiency was monitored using qRT-PCR after 48h.

### Cell Proliferation Assay

OECM-1 cells were seeded in 96-well plates at a density of 3×10^3^ cells/well. Post transfection for 48h, cell viability was assessed using MTT (3-(4,5-Dimethylthiazol-2-yl)-2,5-Diphenyltetrazolium Bromide) (Sigma-Aldrich). 10μl MTT (5 mg/ml) was added into each well followed by 100μl dimethyl sulfoxide (DMSO). Absorbance was measured at 570nm using a microplate reader (Synergy^™^ H1; Agilent BioTek Instruments).

### Cell cycle analysis

OECM-1 cells were seeded in a 6-well plate at the density of 0.5×10^6^ cells/well. 48h posttransfection, cells were harvested and fixed in 1ml of 70% ethanol. For flow cytometric analysis, the fixed cells were resuspended in 1ml of 50μg/ml stock solution of propidium iodide (PI) (Becton Dickinson Biosciences). Further, the samples were incubated for 30 minutes at 4°C and analyzed on a FACSCalibur^™^ Flow Cytometer (Becton Dickinson Biosciences).

### Apoptosis assay

Post transfection, the percentage of apoptotic cells was measured using Annexin V-fluorescein isothiocyanate (FITC)/PI kit using FACSCalibur^™^ Flow Cytometer (Becton Dickinson Biosciences) in accordance with the manufacturer’s instructions (Biolegend).

### Nuclear fragmentation using propidium iodide staining

After 48h of transfection, the monolayered OECM-1 cells were stained with 10μg/ml PI (BD Biosciences) (in PBS) for 5 minutes followed by visualization under the ZOE^™^ Fluorescent Cell Imager at 590nm (Bio-Rad).

### Evaluation of Mitochondrial Membrane Potential (MMP)

Post transfection, the cells were stained with JC-1 (2mg/ml) and incubated for 15 minutes in dark. Changes in MMP were visualized under a fluorescent microscope (Axioskope II, Carl Zeiss).

### Wound healing assay

OECM-1 cells were seeded in a 24-well plate at a density of 0.5×10^5^ cells/well. Post transfection, a scratch was made using a 200μl pipette tip and wounds were photographed at time intervals of 0h, 24h and 48h. The migratory distance of the cells post induction of the wound was measured using ImageJ software (NIH; http://imagej.nih.gov/ij/).

### Transwell invasion assay

In order to assess the invasive capability, OECM-1 cells were seeded in 6-well plates with a density of 0.5×10^6^ cells/well. After transfection, the cells were subjected to a transwell invasion using the QCM Collagen Cell Invasion Assay kit (Millipore) as per the manufacturer’s instructions. 1×10^5^ cells/ml resuspended in serum-free media were added to the top chamber of the inserts whereas 500μl of media containing 10% FBS was added to the bottom chamber. After 48h of incubation, the cells in the bottom chamber were extracted using a detachment buffer. These cells were further stained, lysed and quantified using the 480/520nm filter set in Synergy^™^ H1 microplate reader (Agilent BioTek Instruments).

For imaging, the cells were stained with 10% Giemsa and fixed with formaldehyde after 48h of incubation in the transwell chamber. The cells in the top chamber were removed and those on the bottom of the transwell were visualised at 20x magnification under the ZOE^™^ Fluorescent Cell Imager (Bio-Rad).

### Statistical Analysis

The results of the student’s t-test are represented as mean ± standard deviation (SD) and visualised using GraphPad Prism 6.01 (RRID:SCR_002798) (GraphPad Software, www.graphpad.com). Receiver Operating Curves (ROC) were constructed by plotting relative miRNA expression levels using GraphPad Prism 6.01 software. Principal Component Analysis (PCA) was performed to investigate relationships between samples, and the silhouette score was used to estimate the optimal number of clusters using the prcomp function in the R statistical package. The Association of the miRNA signature with the patient’s overall was evaluated by Kaplan–Meier Survival Curve analysis with the Log-rank statistic using the data available on the TCGA dataset (n=114). Survival analysis was performed using SPSS software (RRID: SCR_002865) (v19.0) (SPSS, Inc.). All experiments were performed in triplicates. Statistical significance was set at p-value<0.05.

## Results

### Identification and characterization of salivary exosomes derived from OSCC patients

The isolated salivary exosomes from OSCC patients were characterized using NTA, TEM, and tetraspanin expression. The NTA results showed a single peak for the concentration of exosomes in the size range of 30-40nm. Salivary exosomes derived from OSCC patients showed a 10-fold higher concentration (7.31 × 10^8^ particles/mL) compared to controls (6.67×10^7^ particles/mL) (Supplementary Fig. S2A). TEM results ( Supplementary Fig. S2B) represent sphere-shaped vesicles with a mean radius of 40-50nm, consistent with the NTA profiles. Flow cytometry analysis indicated the presence of the tetraspanins CD9, CD63, and CD81 (86-88% exosomes expressed these markers) in salivary exosomes from both patients and healthy counterparts (Supplementary Fig. S2C-D), thus confirming that the isolated vesicles comprise of a pure exosomal population.

### Differentially expressed miRNAs enriched in both salivary exosomes and primary tumours from OSCC patients identified by miRNA profiling

Small RNA sequencing was performed to identify differentially expressed miRNAs in tumours and salivary exosomes derived from OSCC patients as compared to age- and sex-matched controls. An average of 25.4 million reads per sample were obtained for all tissue and salivary exosome samples. After pre-processing, adapter sequence trimming and filtering low-quality reads, the mean Q20 values for tissue- and salivary exosome-derived libraries were 97.6% and 93.40%, respectively. Amongst these, an average of 68% (range 61-75%) of the reads could be mapped to hg38 with high confidence.

188 miRNAs were differentially expressed in tumour tissues, 181 miRNAs in leukoplakia samples, 316 miRNAs in salivary tumour exosomes and 69 miRNAs in salivary leukoplakia exosomes compared to those in their normal counterparts. Amongst these, 47 miRNAs were found to be differentially expressed across all sample types and stages, including leukoplakia lesions (Fig. 1A). A comparison of our results with the data from The Cancer Genome Atlas (TCGA) (n=114) database showed that 12 of these miRNAs were commonly differentially expressed between both datasets with a stringent cut-off for adjusted p-value (p<0.01) and log_2_FC > 2 (Fig. 1B). Seven miRNAs (miR-140, miR-143, miR-145, miR-30a, miR-21, let-7i, miR-423) were found to have significant differential expression in both tissue and salivary exosomes derived from OSCC patients compared to controls using qRT-PCR (Fig. 1C). Since the differential expression of these miRNAs is significant in both tissue and salivary exosomes of OSCC patients, we tested their utility as biomarkers for detecting OSCC patients.

**Figure 1:**
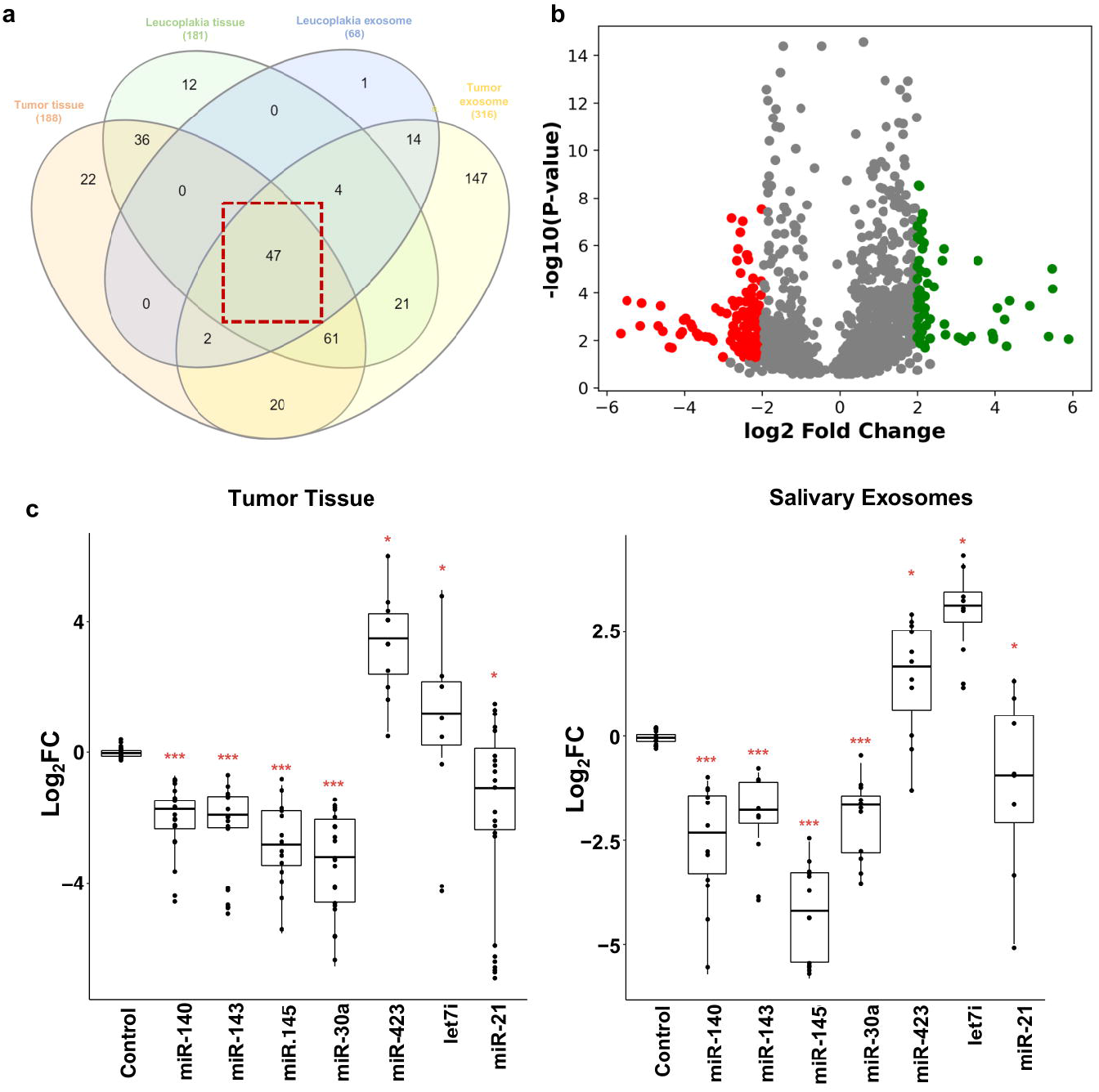
Differential expression of miRNAs observed in tissue and salivary exosomes of OSCC patients. **(a)** Venn diagram representing the forty-seven common miRNAs that were differentially expressed in tissue and salivary exosomes derived from OSCC and leucoplakia patients at nominal adj p < 0.05. **(b)** Volcano plot depicting the twelve significantly expressed miRNAs in both our and TCGA datasets (p < 0.001). **(c)** Box plots depicting the differential expression patterns of seven miRNAs in larger cohorts of tumour tissue and salivary exosomes of OSCC and leucoplakia patients using real-time PCR (***p ≤ 0.001). U6 small RNA was used as internal control, and the relative mRNA levels were analysed using the ddCt method. Error bars represent mean ± SD of three independent experiments.

### A three miRNA signature is able to detect OSCC patients

ROC curves were plotted using the different combinations of these 7 miRNAs to determine their utility in detecting OSCC patients. The combination of miR-140, miR-143, and miR-145 showed an AUC of 0.93 (p < 0.0001) with 93.02% sensitivity and 93.33% specificity in tissue samples compared to the various other combinations of 7 miRNAs (Fig. A). Additionally, salivary exosomes revealed an AUC of 0.99 (p < 0.0001) with 98% sensitivity and 99% specificity using the miR-143/miR-145/miR140 combination (Fig. 2A).

**Figure 2:**
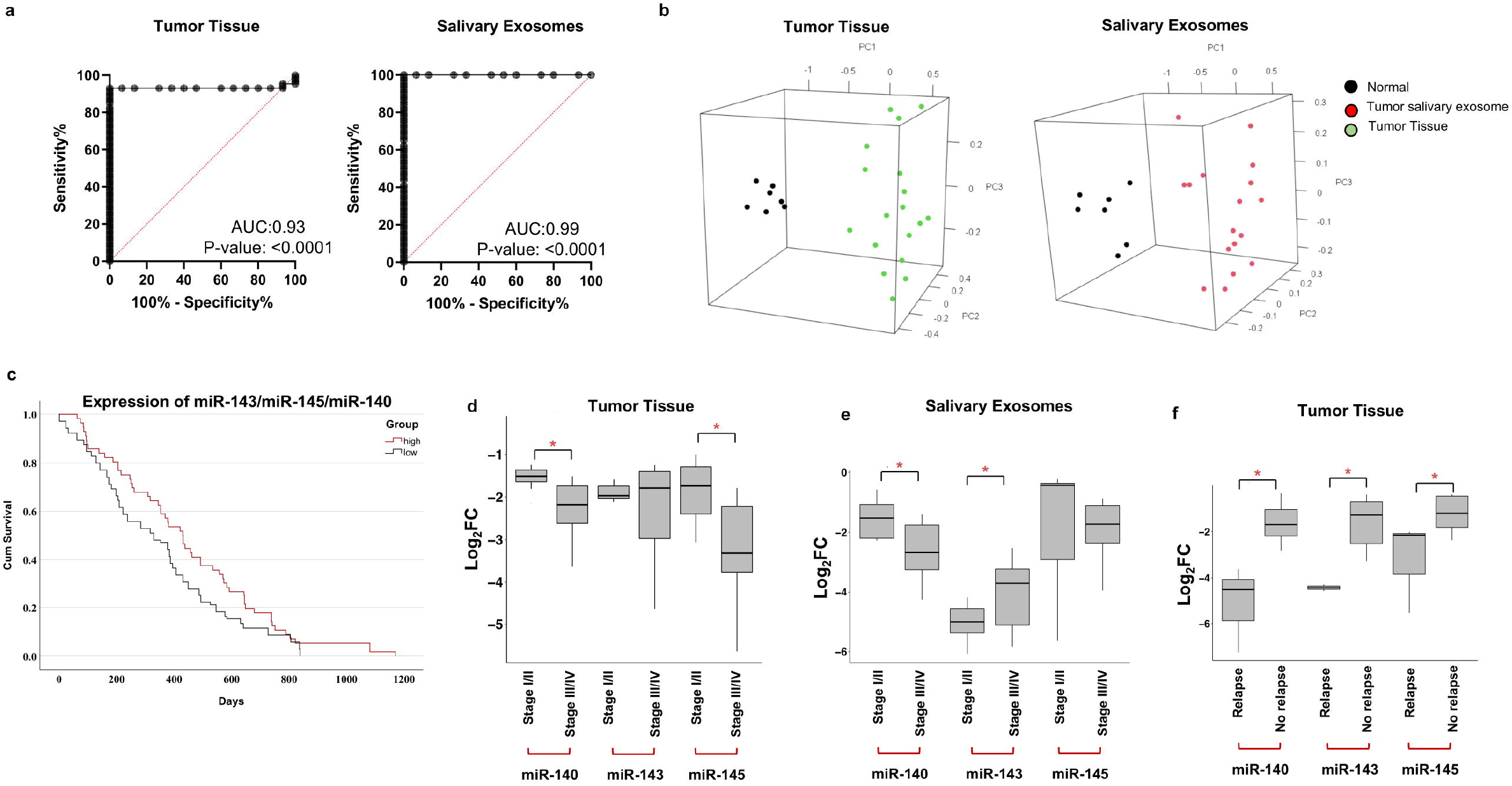
3 miRNA panel can differentiate between cancerous and non-cancerous samples with very good sensitivity and specificity. **(a)** In tissue cohort, the combined measure of sensitivity and specificity of 3 miRNAs (miR-143, miR-145, and miR-140) was represented by AUC of 0.93 (p < 0.0001) with 93.02% sensitivity and 93.33% specificity. In the salivary exosome cohort, the combined AUC of 3 miRNA was 0.99 (p < 0.0001) with 98% sensitivity and 99% specificity. A diagonal reference line acts as a performance measure of the diagnostic test. Note: AUC-area under the curve, CI-confidence interval. **(b)** Representative images are 3D plots of PCA performed using the expression of 3 significantly expressed miRNAs. These plots suggest that expression of these 3 miRNAs can distinguish OSCC tissue samples and salivary exosomes from their normal counterparts with 81.12% variance and 89.98% variance respectively. **(c)** Kaplan–Meier plot of overall survival of patients with high and low expression of miR-143, miR-145, and miR-140 generated from data available on TCGA_HNSC and data obtained from the literature. **(d-f)** Association of identified 3-miRNA signature with various clinicopathological parameters. Representative box plots depict relative expression patterns of miRNAs in early stages (Stage I/II) vs late stages (Stage III/IV) of OSCC patient-derived **(d)** tissue samples and **(e)** salivary exosomal samples. **(f)** Representative box plots depict relative expression patterns of tumour-derived miRNAs in recurrent vs non-recurrent OSCC patients. The expression levels of miRNAs were estimated using real-time PCR. The data were normalised with U6 values, and the relative miRNA levels were analysed using the ddCt method. Error bars represent mean □±□SD of three independent experiments (*p ≤ 0.05; **p ≤ 0.05; ***p ≤ 0.001).

PCA models were constructed for salivary exosomes and tissues derived from OSCC patients as compared to their healthy counterparts, using the expression levels of the 3-miRNA combination. The PCA model was able to segregate discrete clusters of OSCC patients and healthy controls, suggesting the effectiveness of 3-miRNA signature in differentiating cancerous patients from non-cancerous controls (Fig. 2B).

### Three-miRNA signature predicts overall survival and clinically correlates with disease progression and relapse

Survival analysis was conducted based on the expression patterns of miR-143, miR-145, and miR-140 available in TCGA datasets. OSCC patients with low expression of miR-143, miR-145, and miR-140 had a significantly shorter overall survival as compared to those with high expression (p<0.05) (Fig. 2C). On assessing the clinical correlation of the signature we observed a significant downregulation of miR-145/miR-140 and miR-143/miR-140 in the tissue and salivary exosomes of late-stage (stage III and IV; n=30; p<0.05) OSCC patients respectively (Fig. 2D,E). Additionally, miR-143, miR-145, and miR-140 were significantly downregulated (p<0.05) in recurrent OSCC tumours (n=8) as compared to non-recurrent cases (n=8) (Fig. 2F). Collectively these findings suggest that the expression of the 3-miRNA signature can predict poor patient survival, disease progression, and recurrence in OSCC patients.

### Transcriptome sequencing, TCGA datasets and target prediction analysis identifies regulatory miRNA-mRNA networks responsible for OSCC disease progression

To understand the gene regulatory networks modulated by the identified miRNAs, matched samples of OSCC patients were subjected to transcriptome sequencing. RNA-Seq analysis identified 10660 and 11632 differentially expressed mRNAs in tumour and leukoplakia samples of OSCC patients, respectively, compared to their non-cancerous counterparts. On comparing these results with the TCGA datasets, we found a total of 5868 mRNAs that were significantly expressed (log_2_FC>2; adj p-value<0.01) across these datasets irrespective of their disease progression and sequencing platforms.

Target genes of identified miRNA signature were predicted using TargetScan, miRDB and miRWalk software. A total of 315 genes were predicted as potential targets that were differentially expressed in leukoplakia and OSCC patients. On constructing a protein-protein interaction network of the identified genes, three enriched networks (Z-scores: 41, 36, and 35) were generated by IPA (Supplementary Table S2). On merging these networks, we identified top 16 nodes that were further substantiated using the STRING database and CytoHubba plugin in Cytoscape (Supplementary Fig. S3, Supplementary Table S3) (16, 18, 19). Amongst these, five genes viz *HIF1A*, *CDH1*, *CD44*, *EGFR*, and *CCND1* formed dense hubs which are putatively reported to be responsible for OSCC progression (Fig. 3A) (20–27) Subsequently, the differential expression profiles and survival analysis of top 16 hub genes were assessed. A significant upregulation (-log2 (TPM+1)=±2, p< 0.01) was observed in *LOX*, *STAT1*, *PDGFRB*, *JAG1*, *HIF1A*, *GJA1*, *ETS1*, *FLT1* genes in OSCC patients (Supplementary Fig. S4). Moreover, Kaplan-Meier analysis suggested that higher expression of *CD44* (p<0.01), *EGFR* (p < 0.01), *GJA1* (p<0.03), *HNF4A* (p< 0.01), and *CCND1* (p<0.001) significantly correlated with poor overall survival of patients (Supplementary Fig. S5). Further, pathway analysis of the identified miRNA-mRNA networks using IPA (17), identified epithelial-mesenchymal transition (EMT) as the most statistically significant pathway involving a higher number of overlapping genes (Fig. 3B). Collectively, these findings indicate the relevance of identified 3-miRNA signature and its associated gene networks in disease progression and poor prognosis of OSCC patients.

**Figure 3:**
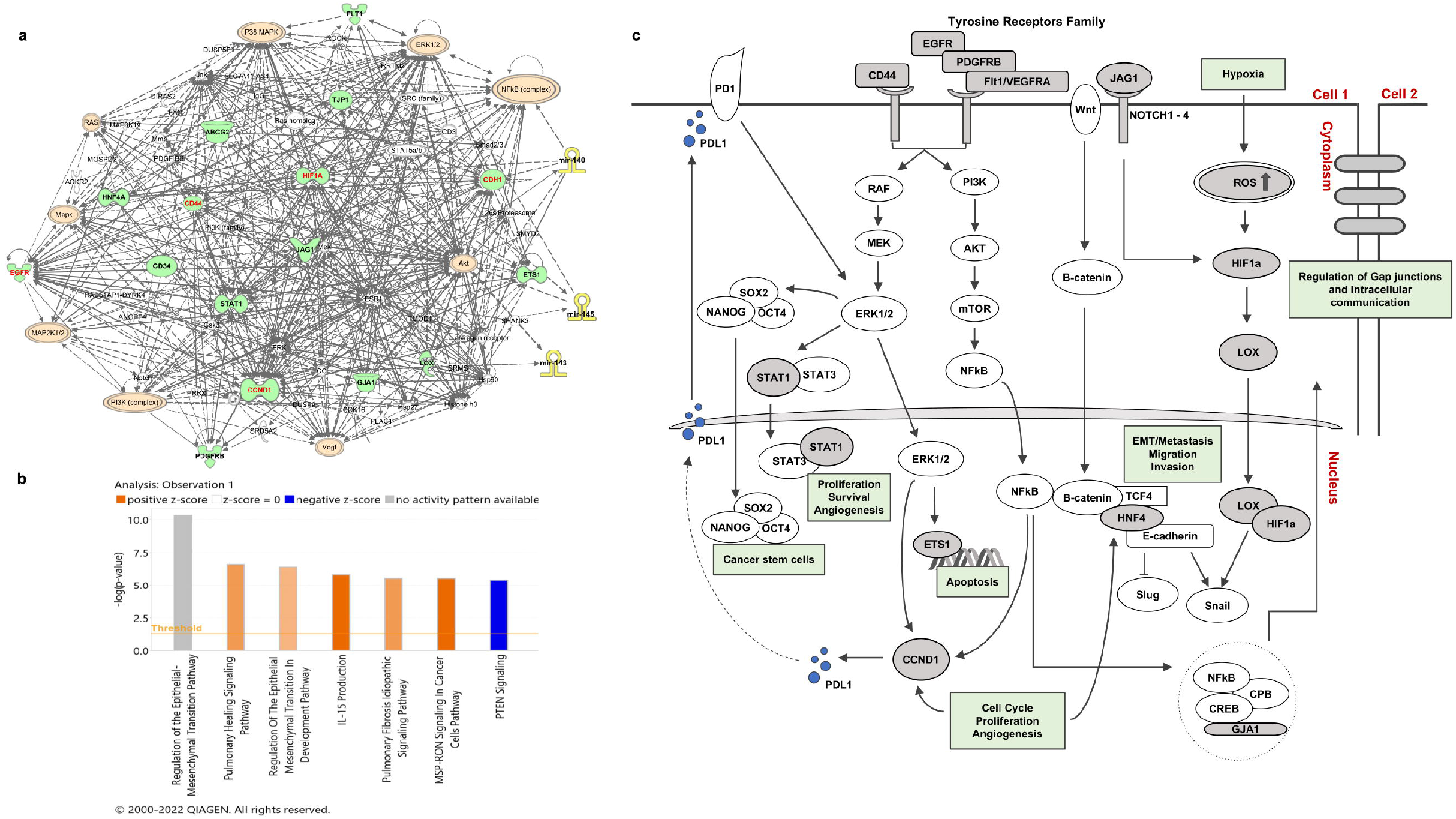
miRNA-mRNA gene network and canonical pathways identified by Ingenuity Pathway Analysis. **(a)** The miRNA-mRNA gene network from experimentally observed miRNA-mRNA interactions of 3-miRNAs and 16 hub genes. Each network displays the genes/gene products as nodes (different shapes representing the functional classes of gene products) and the biological relationships between the nodes as lines. Full lines represent direct interactions and broken lines, indirect interactions. The 3 miRNAs are represented as yellow nodes. Molecules shown in red represent the most centrally located nodes. Orange-colored nodes depict the key molecules responsible for deregulating the major signaling pathways in tumori genesis. **(b)** Differentially expressed canonical pathways targeted by 16 hub genes. Higher −log Benjamini-Hochberg (B-H) p-value corresponds to the more significant pathway. Z score is the activation score for a pathway. The Z score reflects the activation (positive-orange colour) or inhibition (negative-blue colour) of a pathway due to the changes in the expression of genes involved in those pathways or functions. Z-score was not reported for EMT regulation pathway in the IPA libraries **(c)** Schematic representation of putative signalling pathways regulated by the identified miRNA-mRNA networks responsible for OSCC progression. The processes in green boxes are associated with the hallmarks of cancer.

### Overexpression of miR-143/miR-145/miR-140 inhibits cell proliferation and induces G2/M cell cycle arrest

To understand the effect of 3-miRNA signature on functional mechanisms in OSCC, miR-143,miR-145, and miR-140 mimic combination was transfected in OECM-1 cells at 100nM concentration. The transfection efficacy of this mimic combination was measured by realtime PCR. Our results demonstrated a significant upregulation of miR-143, miR-145, and miR-140 by 5.13±0.9 fold, 18.04±2.55 fold, and 23±2.46 fold respectively (Fig. 4A). Consequently, to evaluate the effect of 3-miRNA signature on cell proliferation, MTT assay was performed. A 20% reduction in cell proliferation (p<0.01) compared to scrambled controls was observed 48 hours post-transfection (Fig. 4B).

**Figure 4:**
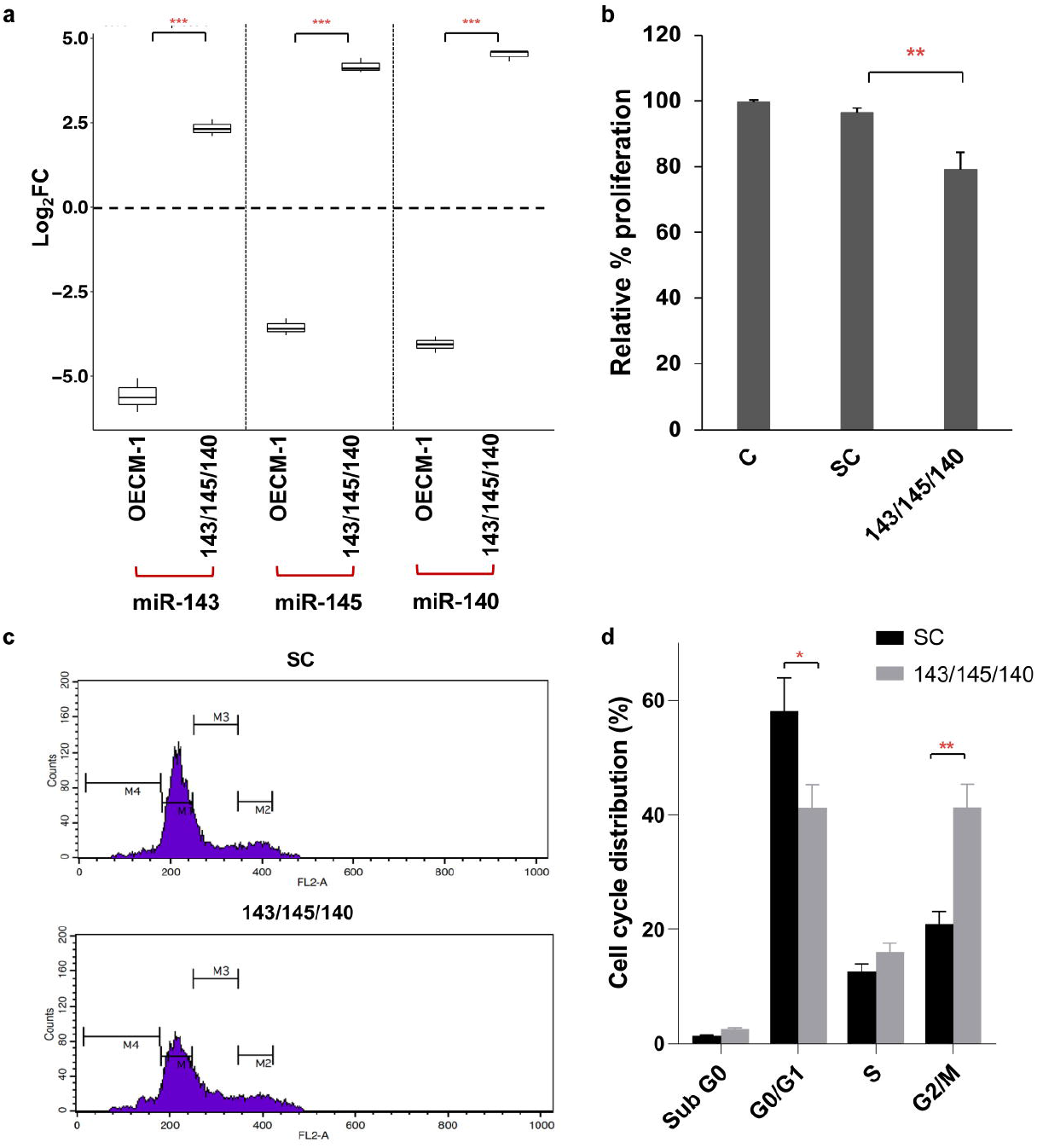
miR-143, miR-145, and miR-140 inhibit OECM-1 cell proliferation and induce a G2/M arrest. **(a)** The expression of miR-143, miR-145, and miR-140 is downregulated in the OECM-1 cell line. After transfecting OECM-1 cells with a combination of miR-143, miR-145, and miR-140 at a total concentration of 100nM for 48 h, a significant increase in the expression of these miRNAs can be seen. U6 small RNA was used as internal control, and the relative miRNA levels were analysed using the ddCt method. Error bars represent mean ± SD of three independent experiments. **(b)** MTT cell proliferation assay was performed after transfecting OECM-1 cells with the 3-miRNA signature. The graph shows the relative cell proliferation rates of untreated controls (C), SC, and transfected (143/145/140) samples. **(c)** Phases of the cell cycle were analysed based on the PI-stained DNA content of OECM-1 cells transfected with miR-143, miR-145, and miR-140 mimics using a flow cytometric analysis. NC miRNA mimic was used as control. The X-axis of the histogram indicates the intensity of the PI signal and the y-axis represents the cell count. **(d)** Representative bar graph shows percentage distribution of cells across cell cycle phases. A G2/M phase arrest can be seen in OECM-1 cells transfected with a combination of miR-143/miR-145/miR-140 mimics. Error bars represent mean □±□SD of three independent experiments (*p<0.05, **p<0.01).

Cell cycle distribution of OECM1 cells transfected with mimics was measured using flow cytometry. Post transfection, a significant increase was observed in the G2/M (41.73±5%) phase compared to the SC (20.93±2.1%; p<0.01). Simultaneously, there was a significant decrease in cells in the G0/G1 phase (41.16±4.8%) compared to SC (58.1±6 %; p<0.05) (Fig. 4C, D). Collectively these results suggest that miR-140, miR-143, and miR-145 restrict the proliferative capacity of tumour cells by inducing G2/M arrest.

### Upregulation of 3-miRNA signature triggers programmed cell death

To elucidate the effect of miR-143/miR-145/miR-140 combination on apoptotic cell death, we performed annexin-V/PI staining 48 □h post-transfection. Upregulation of the 3-miRNA signature resulted in 17.33±3.36% Annexin+/PI-cells compared to 4.33±2.48% Annexin+/PI-cells (p<0.01) in SC, indicating increased apoptosis in the mimic transfected cells (Fig. 5A, B). Subsequently, nuclear staining demonstrated a significant number of PI-stained cells with nuclear fragmentation and chromatin condensation which are prominent characteristics of apoptosis in mimic transfected cells compared to SC (Fig. 5C). Based on these findings, it was necessary to explore the mechanism by which this 3-miRNA signature induced apoptotic cell death. Considering that alteration in the mitochondrial membrane is a crucial event of intrinsic apoptosis, we examined the reduction in mitochondrial membrane potential using JC-1 staining. Our analysis indicated that the transfected cells exhibited a significant reduction in the MMP which was signified by a decrease in red fluorescence and a relative increase in green fluorescence compared to SC (Fig. 5D). Collectively, these findings suggest elevated levels of miR-143/miR-145/miR-140 affects the mitochondrial function and drives oral cancer cells towards mitochondria-mediated apoptosis.

**Figure 5:**
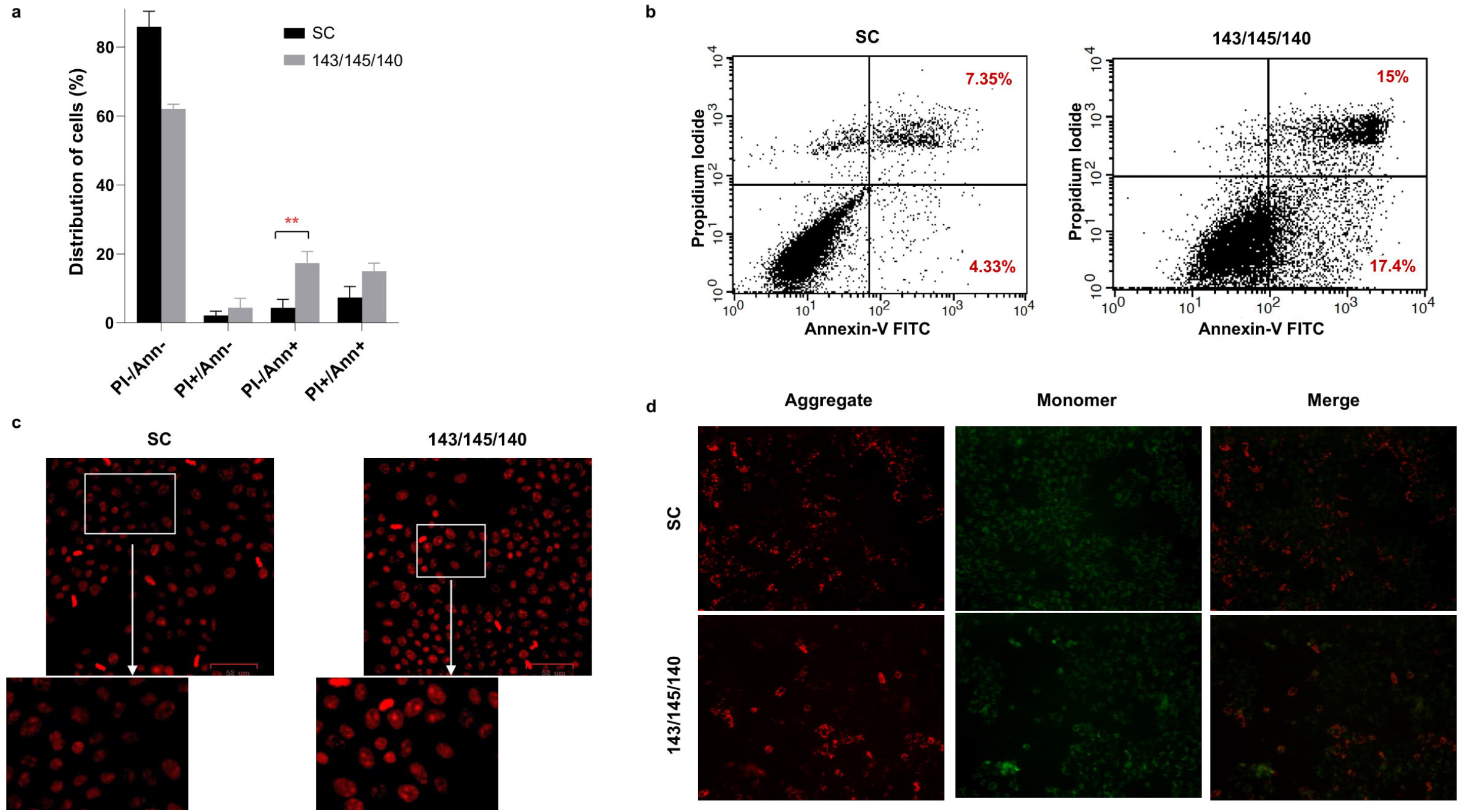
miR-143, miR-145, and miR-140 induce early apoptosis in OECM-1 cells. **(a,b)** Annexin-V/PI staining was used to investigate the apoptotic rates of OECM-1 cells after transfection with SC and miR-143/miR-145/miR-140 mimics. A greater population of early apoptotic cells (Annexin-V-FITC+/PI-) can be seen when OECM-1 cells were transfected with the combination of miRNA mimics (17.33±3.36%) as compared to SC (4.33±2.48%). **(c)** Microscopic images of OECM-1 cells transfected with the 3-miRNa signature and SC. Fragmented nuclei and condensed chromatin can be seen post-transfection of the signature thus confirming the generation of apoptotic bodies.**(d)** Mitochondrial membrane potential was analysed by JC-1 staining. A reduction in the red/green fluorescence ratio indicates a reduction in the mitochondrial membrane potential of cells. This was observed in cells transfected with the combination of miR-143/mir-145/miR-140 mimics. All data are expressed as the mean ± standard deviation. FITC, fluorescein isothiocyanate; PI, propidium iodide. Error bars represent mean □±□SD of three independent experiments (**p ≤ 0.01)

### Enhanced expression of 3-miRNA signature reduces the migratory and invasive potential of OECM-1 cells by reversing epithelial-mesenchymal transition

Given the aggressive nature of oral cancer, we examined the effect of miR-143/miR-145/miR-140 combination on the migratory and invasive potential of OECM-1 cells. On subjecting the cells to wound healing assay, a significant reduction was observed in the migratory ability of mimic transfected cells (230.85±30.56μm) as compared to SC (420.44±35.01μm; p<0.01) (Fig. 6A, B). Additionally, qualitative and quantitative analysis of trans-well assay demonstrated a significant reduction in the invasive potential of mimic transfected OECM-1 cells (42.34±18%) as compared to SC (81.46±6%; p<0.05) (Fig. 6C, D). Hence, these results along with the clinical findings and pathway analysis highlighted the role of miR-143/miR-145/miR-140 in disease progression, aggressiveness and prompted the assessment of its effect on EMT phenomena. Microscopic examination demonstrated an epithelial morphology in a majority of transfected cells as compared to an elongated, mesenchymal-like morphology in SC cells (Fig. 6E), thereby suggesting a reversal of EMT in OECM1 cells. Further, E-cadherin and N-cadherin expression patterns were estimated in OECM-1 cells pre- and post-transfection using qRT-PCR. E-cadherin and N-cadherin expression decreased significantly compared to SC post-transfection with miRNA mimics (Fig. 6F). Therefore, these results show that overexpression of 3-miRNA signature has a vital role in inhibiting cell migration and invasion by inducing mesenchymal-epithelial transition.

**Figure 6:**
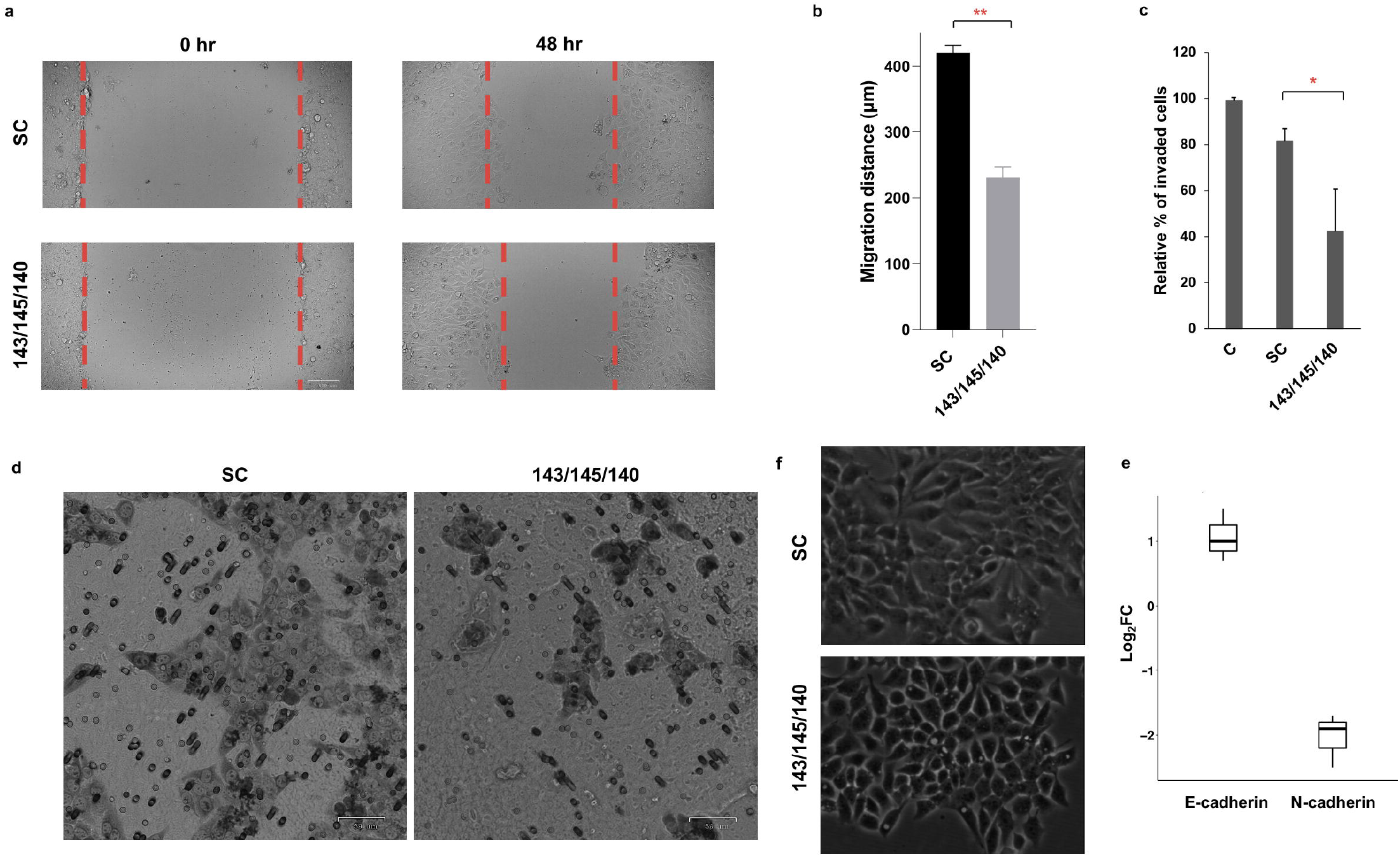
3 miRNA panel inhibits OECM-1 invasion and migration. **(a,b)** Cell migration using a wound healing assay shows a significant reduction in the migratory ability of OECM-1 cells transfected with miR-143/miR-145/miR-140 mimics as compared to those treated with SC. **(c,d)** A transwell assay for cell invasion showed a significant reduction in the proportion of invaded cells when transfected with the combination of 3 miRNA mimics as opposed to SC. (Scale: 59um). **(e)** Phase contrast images show clear visual reduction in the proportion of mesenchymal cells after transfecting OECM-1 cells with the combination of 3 mimics. On the contrary, a greater proportion of mesenchymal cells can be seen in the SC population. **(f)** Representative bar graphs of qRT-PCR-based expression analysis of E-cadherin and N-cadherin in OECM-1 cells after transfection depict relative upregulation of E-cadherin and downregulation of N-cadherin. The expression levels of genes were estimated using real-time PCR. The data were normalised with β-actin values, and the relative gene expression levels were analysed using the ddCt method. Error bars represent mean □±□SD of three independent experiments ( ***p ≤ 0.001).

## Discussion

OSCC has a dismal survival rate owing to late-stage tumour detection making it imperative to identify and develop novel non-invasive, high-screening value tests. The utility of miRNAs as biomarkers of disease progression holds a unique opportunity as they regulate gene transcripts during cellular processes and can serve as indicators of disease status in body fluids. In this study, we conducted a comprehensive analysis of small RNA and transcriptome profiles derived from salivary exosomes and tumour tissue samples from matched OSCC patients. Our small RNA sequencing analysis identified a 3-miRNA signature (miR-143, miR-145, and miR-140) that demonstrated significantly higher sensitivity and specificity in effectively differentiating the cancerous patients from their control counterparts. Further, on assessing the clinical relevance of this 3-miRNA signature, a significant association was observed between suppression of these miRNAs with disease progression, and relapse in oral cancer patients. Thus, these findings reveal the potential role and clinical utility of this 3-miRNA signature in monitoring disease progression, poor prognosis and relapse in oral cancers. In corroboration with our findings, enhanced expression of miR-140 has been reported to significantly correlate with improved overall survival of head and neck cancer patients (28). A study by Bufalino *et al*. has revealed that suppression of tumour-derived miR-143/145 cluster clinically correlated with the progression of the disease, lymph node involvement, and poor survival rates in oral cancer patients (29). Moreover, downregulation of exosomal miR-145 has been previously identified as an important marker for early detection and risk prediction of oral cancer (30). Some of the major confounders of these studies are-small unmatched patient cohorts, the inability to monitor the disease in real-time and the inability to explain the underlying mechanisms responsible for regulating OSCC. To the best of our knowledge, this is one of the preliminary studies that highlight the potential clinical utility of this 3-miRNA signature in predicting disease progression using a liquid biopsy approach.

Given that 60% of protein-coding genes in humans are regulated by miRNAs (6), it is imperative to elucidate the effect of miRNA-mRNA networks on cellular function (7). These miRNA-mRNA interactions and their underlying mechanisms are responsible for the transformation of normal epithelial cells into a malignant phenotype; however, this interplay is completely unexplored in OSCC. Thus, in this study, we performed paired small RNA and transcriptome sequencing to explore the potential of these miRNAs in modulating vital genes responsible for the progression of OSCC. Our integrated network analysis revealed that the identified 3-miRNA signature could potentially target 16 hub genes. Among these, *HIF1α*, *CDH1*, *CD44*, *EGFR*, and *CCND1* were identified as the top hub genes that have also been reported as driver candidates responsible for oral cancer initiation and progression (31–34). Interestingly, miR-143 has been reported to target CD44v3 which has a vital role in inhibiting migration and invasion of oral cancer cells (35). Additionally, Shao et al., demonstrated that miR-145 targeted cyclin D1 expression could effectively suppress OSCC cell growth by regulating cell cycle (36). Collectively, these findings substantiate that miR-143 and miR-145 target a few of the identified genes and have a vital role in OSCC disease progression. In terms of the miRNA-mRNA association, miR-140 has been reported to suppress the invasive and angiogenic properties by altering EGFR levels in lung cancer (37). However, similar interactions between EGFR and miR-140 have not been explored in oral cancers.

Alterations in miRNA-mRNA interactions have an impact on canonical signaling pathways and their controlled feedback mechanism resulting in OSCC disease progression (38–40). Therefore, we explored the downstream signaling pathways that were modulated by the involvement of these miRNA-mRNA networks. Our pathway analysis implied that 3-miRNA signature had the potential to indirectly regulate the EMT pathway by targeting the top 5 hub genes (EGFR, CCND1, CD44, HIF1A and CDH1). EGFR is frequently overexpressed in OSCC and is a primary factor responsible for regulating EMT (41, 42). It has been established that the internalization of EGFR present in OSCC exosomes leads to the transformation of normal epithelial cells to a mesenchymal phenotype (27) CDH1, one of the top hub genes, regulates the process of EMT in oral cancers via multiple pathways including the Wnt/β-catenin signalling cascade (20). Moreover, an increasing number of studies have proven the role of CD44 in regulating EMT in head and neck cancers (HNCs) (21–23). In oral cancer, CD44 governs the invasive and metastatic potential of cells, especially the cancer stem cell sub-population, by modulating the PI3K/Akt/GSK3β pathway (24). Interestingly, it has been reported that proteins of the NOTCH receptor family, one of the most frequently altered pathways in HNCs, increase the recruitment of HIF-1α to the LOX promoter region which in turn activates the SNAIL protein, resulting in migration, invasion, and EMT (25). Furthermore, CCND1 (a gene regulated by the 3 miRNA cluster) has also been reported to induce EMT by regulating its transcriptional factor SNAIL1/2 and modulating the G1/S phase of the cell cycle mechanism (26). Collectively, these observations reveal an inevitable role of the 3-miRNA cluster in modulating the EMT mechanism in oral cancers.

Further, we aimed to investigate the functional role of the 3-miRNA signature using an overexpression model in an oral cancer cell line. Our study demonstrated that overexpression of miR-143, miR-145, and miR-140 reduced cell proliferation, induced G2/M cell cycle arrest and triggered apoptosis via mitochondrial pathway. Our findings corroborated with previous studies that have observed a significant reduction in cell proliferation and apoptosis induced by the overexpression of miR-143 and miR-145 in oral cancers (43, 44). miR-140-5p overexpression was reported to suppress cell proliferation, promote apoptosis, and hamper the progression of cell-cycle in OSCC (45). We also found that overexpression of miR-143, miR-145, and miR-140 induced MET by reducing the invasive and migratory potential in OECM-1 cells. These findings in parallel with pathway analysis, suggest a significant role for the 3-miRNA cluster in transcriptionally regulating genes responsible for the EMT process. The involvement of miR-143, miR-145, and miR-140 individually in regulating the process of EMT and metastasis has been reported in gastric and breast cancers (46–48). However, the association of this 3-miRNA signature with disease progression, aggressiveness, and modulation of EMT has not been explored in oral cancers.

In summary, our findings reveal the diagnostic potential of this 3-miRNA signature using liquid biopsy approach and uncover a plausible mechanism by which dysregulation of these tumour-suppressive miRNAs could effectively convert a normal epithelial phenotype into oral carcinoma.

## Data Availability

The data generated in this study are available upon request from the corresponding authors.

## Figure Legends

**Supplementary Figure S1: Overview of the study design.** This figure explains the design of this study. It shows the sequencing and data analysis pipeline that was followed.

**Supplementary Figure S2: Characterization of salivary exosomes from OSCC patients. (a)** Estimation of concentration and size of salivary exosomes derived from OSCC patients using Nanoparticle Tracking Analysis (NTA). The y-axis represents signal intensity and the x-axis indicates the size distribution of particles in NTA. **(b)** Representative TEM images of exosomes from OSCC patients, (Scale: 200 nm); **(c)** Representative plots of salivary exosomes stained with anti-CD63-Alexa fluor 488, anti-CD81-PE, and anti-CD9-APC antibodies analyzed by flow cytometry.

**Supplementary Figure S3: The gene network of mRNAs constructed using Cytoscape with Maximum Clique Centrality (MCC) Score.** Protein–Protein interaction network generated in Cytoscape using STRING plugin. The hub gene analysis was performed by cytoHubba. The blue-colored nodes represent the 16 nodes with the highest MCC score.

**Supplementary Figure S4**: **Differential expression of top hub genes based on the HNSCC TCGA dataset using the GEPIA platform.** Differential expression of **(a)** LOX **(b)** STAT1 **(c)** PDGFRB **(d)** JAG1 **(e)** HIF1α **(f)** GJA1 **(g)** EST1 **(h)** FLT1 that were found to be significantly expressed in the HNSCC TCGA dataset as compared to their representative controls (p < 0.01), −log2(TPM+1).

**Supplementary Figure S5: Overall survival analysis of hub genes performed using the GEPIA platform.** Overall survival analyses performed using the GEPIA platform where patients with hub genes **(a)** CD34 **(b)** STAT1 **(c)** CD44 **(d)** EGFR **(e)** GJA1 **(f)** HNF4A **(g)** CCND1 expression above the median are indicated by red lines, and black lines indicate patients with hub gene expression below the median. Log-rank p < 0.05 was considered to indicate a statistically significant difference.

## Notes

<u>**Additional Information: Funding:**</u> This work was supported by The Gujarat State Biotechnology Mission-Financial Assistance Programme under the Grant Number: 0P3C9K; and Ahmedabad University Startup grant under the Grant Number AU/SUG/SAS/BLS/2018-19/04-Vivek Tanavde_03.22. Financial Assistantship to SP from the National Postdoctoral Fellowship Program by Department of Science and Technology – Science and Engineering Research Board, Government of India (PDF/2017/001912) is gratefully acknowledged. The Ahmedabad University Assistantship program is appreciated for supporting AP with a doctoral fellowship.

<u>**Competing interests:**</u> The authors declare that they have no competing interests

### Competing Interest Statement

The authors have declared no competing interest.

### Summary of Updates

The updated version of the manuscript includes the findings of the functional assays that were conducted to understand the mechanism of the miRNAs using in-vitro models. These assays also substantiate the findings of the pathway analysis, thus we have revised the manuscript substantially. The new version still retains the data obtained from transcriptome and small RNA sequencing followed by qRT-PCR-based validation. Figures 1, 2, 7 and 8 from the previous version have been moved to supplementary data (Supplementary Figures 1, 2, 4, and 5 respectively). In the new version, Figures 1-6 have been revised and updated to incorporate the in-vitro findings. The methodology section has been updated to include additional experiments that were performed. Similarly, the results and discussion sections have also been updated to include additional findings and substantiate the conclusion of the previous version.

