## Supplementary Tables for "A novel 3-miRNA network regulates tumour progression in oral squamous cell carcinoma"

**Supplementary Table 1: Primer sequences of miRNAs and genes.**

| **miRNA** | **Primer Sequence** |
| --- | --- |
| miR-140-5p | **F** 5’-GAGTGTCAGTGGTTTTACCCT-3’ **R** 5’GCAGGGTCCGAGGTATTC-3’ |
| miR-143-5p | **F** 5’-GGGACAGACACCCGTTTTGA-3’  **R** 5’ GTGTTGCCCACGGTAATGCT-3’ |
| miR-145-5p | **F** 5’-CAGAGTGCGTGTCGTGGAGT-3’  **R** 5’-AGGTCCAGTTTTCCCAGG-3’ |
| miR-30a-5p | **F** 5’-GGGCCTGTAAACATCCTCG-3’  **R** 5’-GAATACCTCGGACCCTGC-3’ |
| miR-423-5p | **F** 5’-TTGGAGTAGGTCATTGGGTGG-3’  **R** 5’-CCAAGACATGGAGGAGCCAT-3’ |
| let-7i-5p | **F** 5'-TGAGGTAGTAGTTTGTGCTGTT-3'  **R** 5'-GCGAGCACAGAATTAATACGAC-3' |
| miR-21-5p | **F** 5'-TTTTGTTTTGCTTGGGAGGA-3'  **R** 5′-AGCAGACAGTCAGGCAGGAT-3′ |
| U6 | **F** 5’-CTCGCTTCGGCAGCACA-3’  **R** 5’-AACGCTTCACGAATTTGCGT-3’ |

| **Gene** | **Primer Sequence** |
| --- | --- |
| E-Cadherin | **F** 5’-ATTCTGATTCTGCTGCTCTTG-3’ **R** 5’-AGTCCTGGTCCTCTTCTCC-3’ |
| N-Cadherin | **F** 5’-CCACGCCGAGCCCCAGTATC-3’  **R** 5’-CCCCCAGTCGTTCAGGTAATCA-3’ |
| β-Actin | **F** 5’-CATGTACGTTGCTATCCAGGC-3’  **R** 5’-CTCCTTAATGTCACGCACGAT-3’ |

**Supplementary Table 2**: **The top enriched networks generated by IPA comprising of different sets of differentially expressed genes.**

| **ID** | **Molecules in Network** | **Score** | **Focus Molecules** | **Top Diseases and Functions** |
| --- | --- | --- | --- | --- |
| 1 | ADAM9,CALU,CDH1,CITED2,CTTNBP2NL,DAB2,DLGAP1,DPP4,ERK1/2,FMNL2,GJA1,GRB10,Growthhormone,LMNB2,MMD,Mmp,MYO6,NETO2,NRIP1,NUAK1,PAXBP1,PDE4D,PODXL,PP2A,PRKAA,PTGFR,S1PR1,SET,Shc,SPTBN1,STAT5a/b,STK38L,STRN,TJP1,TRPM7 | 43 | 28 | [Cancer, Organismal Injury and Abnormalities, Respiratory Disease] |
| 2 | ABCE1,ABCG2,ACSL4,AHI1,ANKHD1/ANKHD1-EIF4EBP3, B4GALT6, CCND1, CDK4/6,CDK6,COL4A1,DTL,E2f,EGFR, estrogen receptor, G3BP1, GRHL1, Hdac,JAG1,KLF9,KLHL24,LRP6,MAP2K1/2,Mek,MGAT1,PITX2,PTPRK,RAD18,RAD51,Raf,SACS,SMAD1,Smad2/3,SRGAP1,TEAD1,THNSL1 | 41 | 27 | [Cellular Response to Therapeutics, Embryonic Development, Organismal Development] |
| 3 | 143,ABL1,B4GALT5,BCL11B,BCR,CAMK2G,CARD10,caspase,CD44,Creb,ETS1,FSCN1,GLI3,LAMC1,Mapk,MMP13,MMP3,MYL12A,NFkB (complex),PDGFRB,Pka,Pka catalytic subunit,Pkc(s),PPP3CB,PRLR,RAS,RNF216,ROCK,ROCK1,RTKN,SRGAP3,ST3GAL5,TAX1BP1,THY1,WASF1 | 36 | 25 | [Cancer, Cellular Movement, Gastrointestinal Disease] |
| 4 | AARS1,Actin,BCR (complex),CCNB1,CDCA3,DGKH,E2F3,EGLN,ELK4,ERK,FLNB,Gamma tubulin,Histone h3,HTATSF1,IgG,Jnk,KAT2B,MAP3K4,MTORC1,NECTIN1,P-TEFb,PAN2,PDGF BB,PGM1,PPP1CC,Rb,RNA polymerase II,SLC38A2,SMAD5,STMN1,TCF3,Tgf beta,TM4SF18,TMOD3,ZNF337 | 28 | 21 | [Cancer, Hematological Disease, Immunological Disease] |
| 5 | AKAP12,Akt,Alphatubulin,ANGPT2,APOL4,B4GAT1,CD3,DLC1,DPYSL2,EEA1,F Actin,FLT1,Focal-adhesion kinase,Gsk3,HIF1A,Hsp27,Hsp70,Hsp90,KANK2,LOX,MAP3K5,NUCKS1,P38 MAPK,PAPSS2,PI3K(complex),PI3K (family),PTGS1,RALGDS,Ras homolog,SLC4A7,SLC7A1,SRC (family),STAT1,TUBGCP3,Vegf | 26 | 20 | [Cancer, Cardiovascular System Development and Function, Tissue Morphology] |
| 6 | BCL9,BICC1,BTF3,CDON,CSNK1D,CTNNB1,CUL1,CXCL14,CYP51A1,EID1,EXT1,FER,FGD4,GPR161,GPS1,HAMP,HIC1,HMG CoA synthase,HMGB3,IDI1,KIF3A,LRP4,LZTS2,MARK1,MDFIC,NUCKS1,OSBPL1A,PLPPR4,PYGO1,PYGO2,RGS3,SEMA6A,SFRP4,SOST,TMEM47 | 17 | 15 | [Embryonic Development, Organ Development, Organismal Development] |
| 7 | ACP2,AHR,B4GAT1,CPM,CPSF6,DHRS9,EBF1,EFNB3,ERBB2,EXT2,FCHO2,FUT1,HECW2,HS3ST1,Igh(family),ITGB1,LMBRD2,MED1,NANOG,NUDT21,PHACTR2,PPARGC1B,S100A6,SCARA3,SEL1L3,SEMA7A,SLC43A3,SLC7A8,SLCO4A1,SMARCA4,SYVN1,TMEM106B,TPM4,TSPAN13,TUSC3 | 17 | 15 | [Embryonic Development, Organ Development, Organismal Development] |
| 8 | APH1B,ATG14,BECN1,CERS6,CLSPN,CPEB3,DUT,E2F1,ERK1/2,ESCO2,ESYT1,GTPBP4,HERC2,ISCU,let7,MCM2,NAP1L1,NRBF2,OR51E1,PABPC4,PCSK5,PCSK7,PIK3R4,PSAT1,RPTOR,SNRPC,TAGLN2,THAP12,TP53,ULK2,UVRAG,VEGFD,VRK2,ZFYVE1,ZNF512B | 14 | 13 | [Carbohydrate Metabolism, Lipid Metabolism, Small Molecule Biochemistry] |
| 9 | ANGEL1,ANO6,EIF4E,EIF4EBP2,FLVCR1,HILPDA,HOXA5,HOXB5,IPO7,MXD1,MXRA5,MYC,Nc2,NELL2,NFIB,NMNAT1,NPM1,NUP50,NUP98DDX10,NUPR1,PARP1,PEG10,PLOD1,PLXND1,RPRD1B,SNHG17,SUPT5H,TCAF2,TEP1,USP36,VEGFA,YTHDC1,ZBTB34,ZFP36L1,ZNF521 | 14 | 13 | [Embryonic Development, Organismal Development, Skeletal and Muscular System Development and Function] |
| 10 | ALPK3,APPL1,ARID1A,ARID4A,CEACAM6,CELSR2,CITED1,CLIC3,ESR1,FAM13C,FGFBP1,GOLM1,GPAM,GREB1,H4C3,HDAC1,HECTD1,KRT16,KRT4,KRT6A,KRT6B,MIER1,Mta,PDZK1,PIN1,PKNOX2,PRXL2A,PTGFRN,RNF181,SAP30,SLC2A4,SLC7A2,SNX24,TFF2,TOP2B | 11 | 11 | [Dermatological Diseases and Conditions, Developmental Disorder, Hereditary Disorder] |

**Supplementary Table 3: List of top 16 hub genes with their respective interaction scores generated by the MCC method.**

| **Gene** | **Score** |
| --- | --- |
| *HIF1A* | 75187 |
| *CDH1* | 74301 |
| *CD44* | 74210 |
| *EGFR* | 73498 |
| *CCND1* | 73068 |
| *JAG1* | 35528 |
| *CD34* | 32902 |
| *FLT1* | 25682 |
| *PDGFRB* | 22019 |
| *GJA1* | 16607 |
| *ETS1* | 15977 |
| *STAT1* | 15975 |
| *ABCG2* | 15843 |
| *HNF4A* | 15369 |
| *TJP1* | 15262 |
| *LOX* | 11123 |
| *PDGFRB* | 22019 |
