## Supplementary figures and images for "A novel 3-miRNA network regulates tumour progression in oral squamous cell carcinoma"

### Supplementary Figure 1

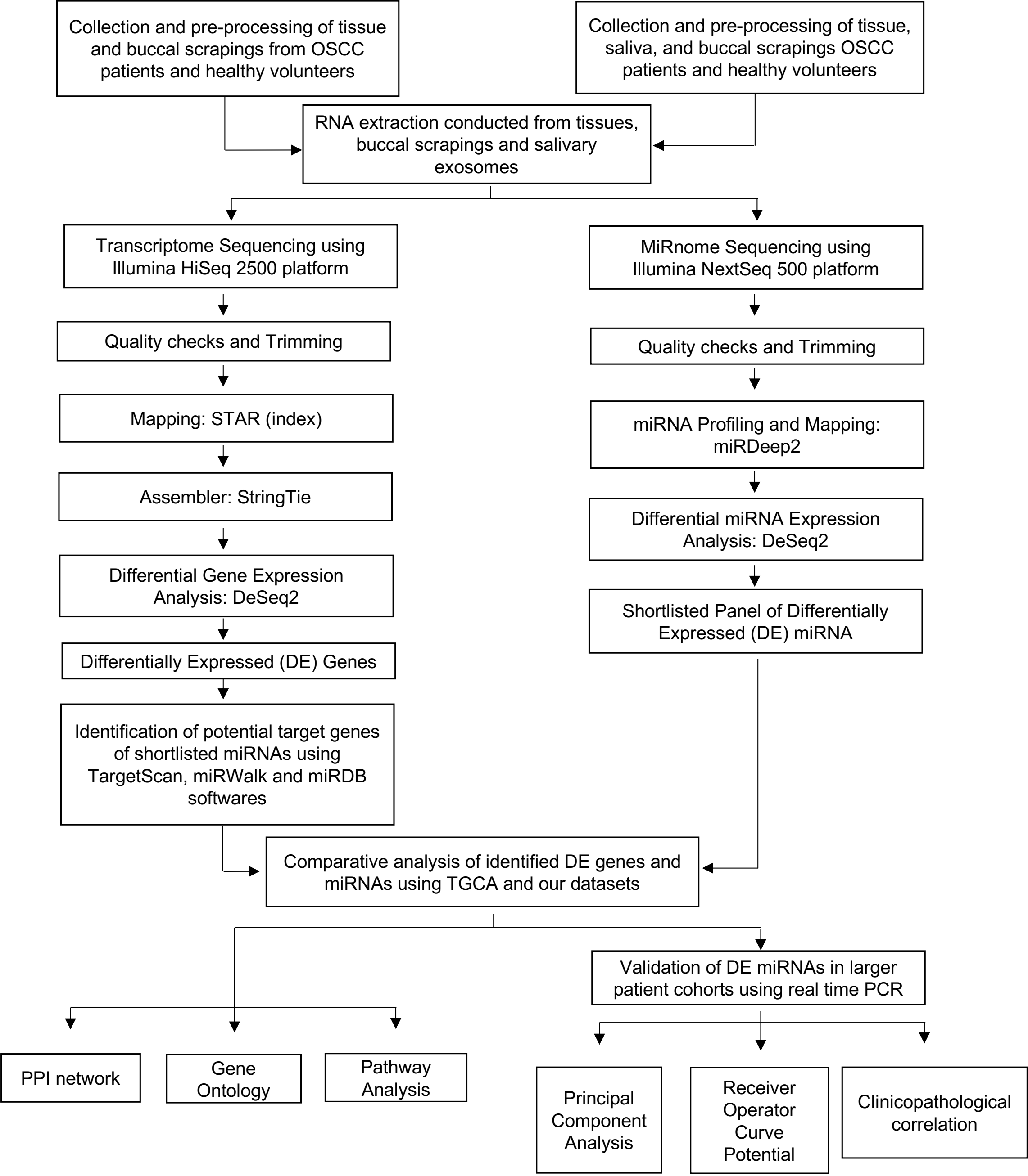

### Supplementary Figure 2

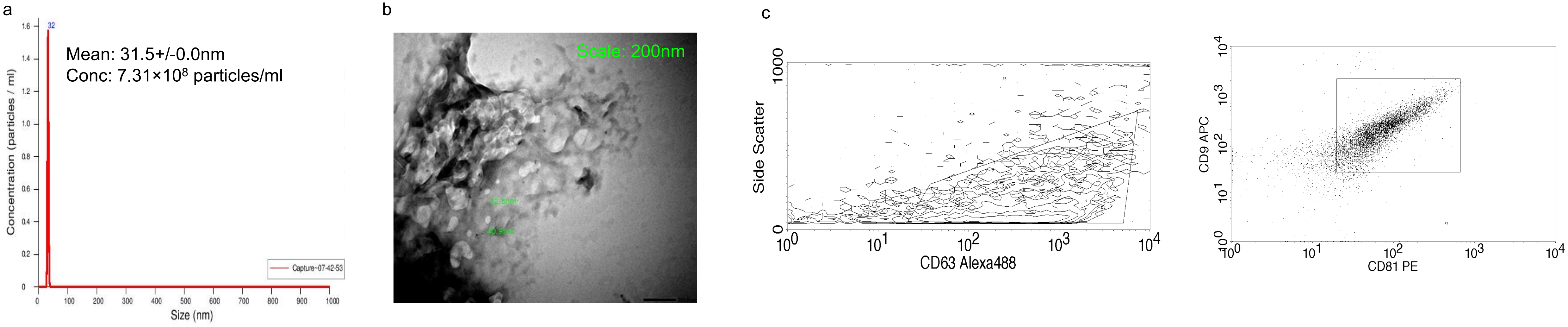

### Supplementary Figure 3

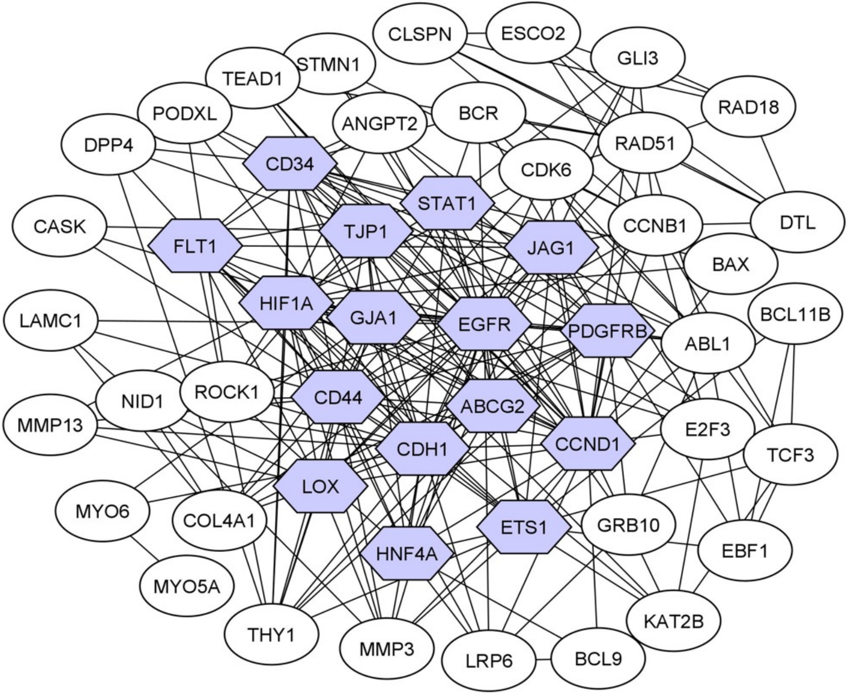

### Supplementary Figure 4

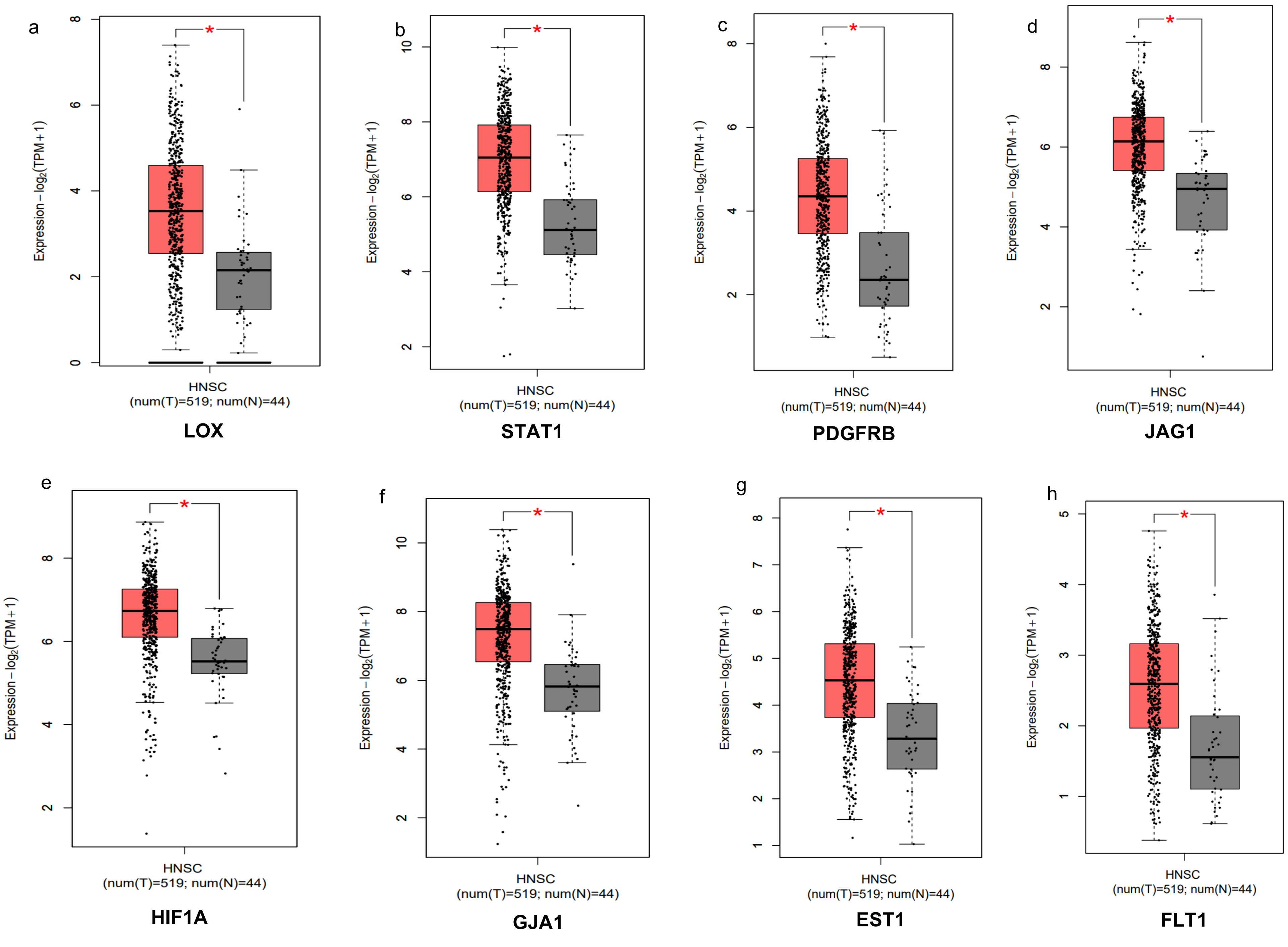

### Supplementary Figure 5

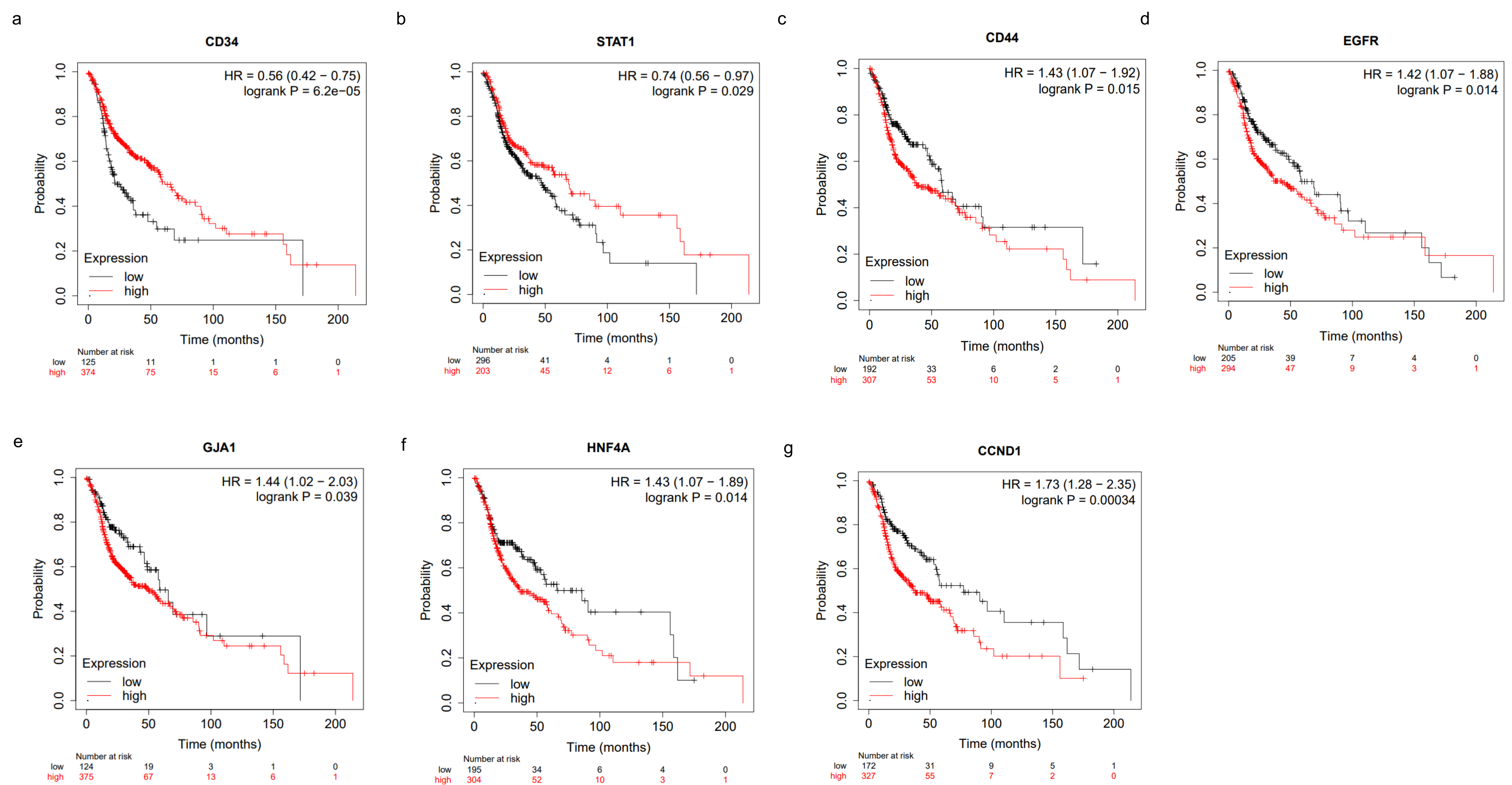
